# Phyllosphere Community Assembly and Response to Drought Stress on Common Tropical and Temperate Forage Grasses

**DOI:** 10.1101/2021.05.10.443535

**Authors:** Emily K. Bechtold, Stephanie Ryan, Sarah E. Moughan, Ravi Ranjan, Klaus Nüsslein

## Abstract

Grasslands represent a critical ecosystem important for global food production, soil carbon storage, and water regulation. Current intensification and expansion practices add to the degradation of grasslands and dramatically increase greenhouse gas emissions and pollution. Thus, new ways to sustain and improve their productivity are needed. Research efforts focus on the plant-leaf microbiome, or phyllosphere, because its microbial members impact ecosystem function by influencing pathogen resistance, plant hormone production, and nutrient availability through processes including nitrogen fixation. However, little is known about grassland phyllospheres and their response to environmental stress. In this study, globally dominant temperate and tropical forage grass species were grown in a greenhouse under current climate conditions and drought conditions that mimic future climate predictions to understand if (i) plant host taxa influence microbial community assembly, (ii) microbial communities respond to drought stress, and (iii) phyllosphere community changes correlate to changes in plant host traits and stress-response strategies. Community analysis using high resolution sequencing revealed *Gammaproteobacteria* as the dominant bacterial class, which increased under severe drought stress on both temperate and tropical grasses while overall bacterial community diversity declined. Bacterial community diversity, structure, and response to drought were significantly different between grass species. This community dependence on plant host species correlated with differences in grass species traits, which became more defined under drought stress conditions, suggesting symbiotic evolutionary relationships between plant hosts and their associated microbial community. Further understanding these strategies and the functions microbes provide to plants will help us utilize microbes to promote agricultural and ecosystem productivity in the future.

**IMPORTANCE:** Globally important grassland ecosystems are at risk of degradation due to poor management practices compounded by predicted increases in severity and duration of drought over the next century. Finding new ways to support grassland productivity is critical to maintaining their ecological and agricultural benefits. Discerning how grassland microbial communities change in response to climate stress will help us understand how plant-microbe relationships may be useful to sustainably support grasslands in the future. In this study, phyllosphere community diversity and composition was significantly altered under drought conditions. The significance of our research is demonstrating how severe climate stress reduces bacterial community diversity, which previously was directly associated with decreased plant productivity. These findings guide future questions about functional plant-microbe interactions under stress conditions, greatly enhancing our understanding of how bacteria can increase food security by promoting grassland growth and resilience.

## INTRODUCTION

Agricultural production, through crops and grazing, is key to global food security. Despite current advances, the number of undernourished people is increasing (1). Grasslands are an important ecosystem with potential to help promote food security through their role in ruminant milk and meat production and reduce greenhouse gas levels by sequestering and storing carbon. Current intensification and expansion practices have taken a serious toll on the environment, causing degradation of land, water, and biodiversity while increasing greenhouse gas emissions (2–4). As global temperatures continue to rise, weather patterns are predicted to drastically change increasing intensity, duration, and frequency of drought in important grassland areas over the next century (5–8). This will not only affect agricultural production, but also increase levels of atmospheric CO_2_ further destabilizing current weather patterns (9). Thus, finding ways to promote grassland health and production in sustainable ways is essential for increasing food security and reducing environmental strain.

The phyllosphere, or aerial surface of plants, is estimated to cover over 10^9^ km^2^ and contain 10^26^ bacterial cells, making it one of the largest microbial habitats on earth (10). Microbes in the phyllosphere promote host fitness through phytohormone and nutrient production, increased stress tolerance, and protection against pathogens (11–13). As a result, phyllosphere communities can support ecosystem functioning (10, 13–16). In order to leverage plant-microbe relationships to promote ecosystem health and agricultural production (17–19), we need a better understanding of which microbes are associated with plant hosts, what drives their community structure, how they support their host, and how microbial communities are changing as a result of climate stress.

Due to temperature fluctuations, changing precipitation patterns, and UV exposure, the phyllosphere is an extreme habitat for microbes. Despite these harsh conditions, phyllosphere communities exhibit strong seasonal and temporal patterns and according to the literature are consistently dominated by *Gammaproteobacteria* and *Alphaproteobacteria* (18, 20, 21). Within *Alphaproteobacteria*, members from the genera *Methylobacterium* and *Sphingomonas* are ubiquitous across many phyllosphere studies. They are hypothesized to be generalists able to survive off of a low abundance of many different substrates making them ideal members of the continuously changing phyllosphere ecosystem (18, 22, 23). Although studies of grassland and tree phyllospheres identified host species as an important driver of community structure (24–27), a recent study found that the bacteria species detected in the phyllosphere were shared across the different host species studied suggesting that phyllosphere communities are composed of generalist bacteria capable of surviving variable environmental conditions (28). These differing trends suggest community assembly is not a completely stochastic process, and what drives assembly is not well understood (19, 21). Because plant-health is reliant on microbial communities, understanding the ecological impacts of climate change on plant-microbe relationships is important to develop strategies that counteract the associated negative effects (19, 29).

In spite of the ecological importance of grasses, few studies have used culture-independent methods to investigate bacterial phyllosphere communities on grasslands. Studies suggest soil acts as a reservoir for phyllosphere microbes and that strong seasonal, environmental, and temporal patterns influence phyllosphere community structure (18, 30). Despite grassland phyllosphere communities showing significant shifts over time, community assembly is also impacted by host identity (27, 30). Experiments with grasses at elevated temperatures showed significant shifts in phyllosphere communities with plant beneficial bacteria decreasing and potential plant and human pathogens increasing (23, 31). Additionally, grass phyllosphere communities are greatly impacted by urbanization (32). More targeted studies are needed to understand how continued climate stress will impact grassland phyllosphere community dynamics.

Grasslands around the world have a wide range of physiological and morphological features including different carbon fixation mechanisms (33). Many tropical grass species utilize the C_4_-pathway, with a 50% improved photosynthetic efficiency over the C_3_-pathway utilized by temperate grass species (34). These physiological differences result in different levels of stress tolerance and response strategies. For example, C_4_ plants have lower stomatal conductance and perform photosynthesis while stomata are closed, resulting in lower transpiration rates and continued biomass production under drought stress (35). Understanding if differences in plant physiology relate to differences in phyllosphere community structure under changing climate conditions will shed light onto drivers of microbial community assembly and determine if universal strategies using microbes to promote grassland health can be developed. Three closely related species of C_4_ tropical grasses and two species of C_3_ temperate grasses were grown under optimal temperature conditions in greenhouses and subjected to different watering regimes to understand (i) how plant host identity influences microbial community assembly, (ii) how microbial communities respond to drought stress, and (iii) if the changes correlate to changes in plant host traits and plant stress response strategies. We hypothesized that (1) each species of plant has distinct phyllosphere communities and more closely related plant species have more similar communities. (2) Drought results in decreased microbial community diversity and changes in microbial community structure on all plant species. Additionally, (3) bacterial communities from more drought tolerant species show less change due to their ability to better grow and survive under drought conditions.

## MATERIALS AND METHODS

Three closely related C_4_ tropical forage grass species (*Brachiaria brizantha*, *Brachiaria decumbens*, and Brachiaria hybrid cv Cobra (CIAT 1794)) and two C_3_ temperate forage grass species (*Festuca arundinacea* and *Dactylis glomerata*), were grown under three different watering conditions: well-watered control, mild drought (MD), and severe drought (SD). Grass species were chosen because they are widely used in forage systems in either tropical or temperate climates. Experiments were set up in the College of Natural Sciences Research and Education Greenhouse at the University of Massachusetts-Amherst. Grass host species and their relatedness were compared to understand if they influenced bacterial community diversity and structure, and to determine if different plant responses to drought stress influence changes in microbial community structure. At each sampling time, mature, but not senescing, leaves were randomly sampled. Drought severity was standardized based on the leaf relative water content (Supplemental Figure S1B and S2A).

### Plant Growth Conditions

#### Temperate Grasses

Endophyte free *Dactylis glomerata* (Orchardgrass) and *Festuca arundinacea* (Tall Fescue) seeds were acquired from Albert Lea Seed Company (Albert Lea, MN, USA). Seeds were germinated in Pro-mix commercial potting media (Quakertown, PA, USA) in October 2018. After 6 weeks, each individual plant was transplanted into its own plastic pot (13 cm diameter by 23 cm height) so that each pot contained a plant from one seed. Pots were filled with soil collected from natural grass fields in Amherst, MA. The top 15 cm of topsoil was collected, all rocks and roots were removed, and soil was immediately transported to the greenhouse for planting. Soil nutrients were tested at the University of Massachusetts Soil and Plant Nutrient Testing Laboratory (Supplemental Table S1). Plants were maintained under greenhouse conditions at 21°C for 16 hours light provided by 1000W metal Halide and 50% 600W High pressure sodium lights and at 18°C for 8 hours during the nighttime. Humidity was set at 50%. Drought experiments began once plants reached 5 months of growth.

#### Tropical Grasses

*Brachiaria brizantha* (CIAT 26564)*, Brachiaria decumbens* (CIAT 6370), and *Brachiaria* hybrid (CIAT 1794) seeds were acquired from CIAT (Cali, Columbia). Seeds underwent acid scarification before germination in Pro-mix commercial potting medium in January 2018. After 4 months, individual plants were transplanted into plastic pots (13 cm diameter by 23 cm height) so that each pot contained one plant. Pots were filled with soil designed to mimic tropical soil. Soil from a forest in Amherst, MA underwent nutrient testing (Supplemental Table S1) to determine nutrient similarity to tropical soil. Soil was collected, rocks and roots were removed, soil was then amended with kaolinite in order to replicate the high clay content found in Amazonian soil, and then soil was transported to the greenhouse for immediate planting. Plants were maintained under greenhouse conditions at 30°C for 16 hours light provided by 1000W metal Halide and 50% 600W High pressure sodium lights and at 26°C for 8 hours during the nighttime. Humidity was set at 60%. Drought experiments began 5 months after seeds were planted.

### Drought Experiment

Fully grown grasses were divided into a control group and two independent drought treatment groups. Control plants were watered to maintain field capacity, MD at 30-50% field capacity, and SD at 10-20% field capacity. Plants were organized in a randomized block design. Field capacity was determined before the start of the experiment by flooding the soil with water, allowing it to drain for 24 hours and then measuring soil moisture with a soil moisture meter. Water used throughout the experiment was standard tap water used in the greenhouse facility.

In temperate grass experiments, each of the three watering treatments consisted of 5 biological replicates for each of the two plant host species, except for the Tall Fescue control treatment which had 4 replicates, for a total of 29 individual plants. For 36 days, the soil moisture content was measured every three days using a MiniTrase TDR with Buriable probe (Soilmoisture Equipment Corp., Goleta, CA, USA). In tropical grass experiments, each of the three watering treatments consisted of 3 biological replicates for each of the three plant host species totaling 27 individual plants. Over 22 days, soil moisture content was measured weekly using an Extech Soil Moisture Meter (Extech, Waltham, MA, USA). The amount of supplemental water was determined based on soil moisture readings for each pot. Because individual plants differed slightly in total biomass, the rate of transpiration was different between each plant and therefore each plant required varying amounts of water to maintain equal soil moisture.

### Plant Health Measurements

Throughout each drought experiment, several plant health measurements were used to assess plant response to drought. These measurements were taken from the same plants from which phyllosphere communities were collected in order to directly correlate plant traits with microbial communities. Leaf relative water content of each grass species was measured weekly (36). Leaf chlorophyll concentration was determined weekly by extracting chlorophyll from the leaves using dimethyl sulfoxide and measuring spectrophotometrically (37). Additionally, leaf cellular membrane stability was determined by measuring electrolyte leakage in temperate grasses (38) three times throughout the experiments on days coinciding with leaf sampling. Leaf relative water content, chlorophyll concentration, and membrane stability were measured in the laboratory after aseptically removing three mature but not senescent whole leaves from each plant. Photosynthetic capabilities and efficiency were further assessed weekly in tropical grasses using a chlorophyll fluorometer (Opti-Sciences Inc., Hudson, NH, USA) following established protocols (39).

Plant growth was non-destructively determined by measuring leaf width on a weekly basis. At the end of the experiment, biomass was measured by harvesting the plants and dividing them into five categories: stems, dead material, roots, mature leaves, and young leaves. Fresh mass was taken for each sample, then dried in an incubator at 70°C for 5 days after which dry mass was determined.

### Bacterial Community Analysis

Phyllosphere communities collected from well-watered control and drought-exposed plants were characterized using 16S rRNA marker gene sequences. In the tropical drought experiment, leaves from each plant were collected for total cell counts and DNA extraction on days 1, 12, and 22, totaling 81 samples. In the temperate drought experiment, leaves were collected on days 1, 14, 23, and 36, totaling 112 samples (one Tall Fescue and one Orchardgrass control sample were removed due to pest infestation).

#### Bacterial Cell Counts

Three leaves (∼1-2 grams) were placed in 50 ml conical tubes with 10 ml of phosphate-buffered saline (PBS), incubated at room temperature for 1 hour, then vortexed horizontally (vortex-adaptor, Qiagen, Germantown, MD, USA) at full speed for 10 min. The PBS leaf wash was collected and the process repeated. Samples were then fixed with 3.7% paraformaldehyde and stored at 4°C. Samples were filtered onto black polycarbonate membrane filters (pore size 0.2 µm, 25 mm diameter) (Steriltech Corporation, Kent, WA, USA), stained using 0.1% acridine orange for 3 min, and analyzed with epifluorescence microscopy by counting 20 fields using SimplePCI (Hamamatsu, Japan) (40, 41).

#### DNA Extraction and Sequencing

Whole leaves were aseptically removed from plants in the greenhouse and stored at 4°C until bacterial community extraction was carried out the same day. For temperate drought, bacterial DNA was extracted using the Nucleospin Plant II Extraction Kit (Machery-Nagel, Düren, Germany) with 3 modifications. Three whole leaves (∼1-2 g) were placed into 15 ml conicals containing 1ml of NucleoSpin Type-B beads and 1.6 ml of Buffer PL1. Tubes were vortexed horizontally for 5 min at room temperature. Aydogan et al. (2018) found that vortexing whole leaf samples in tubes with lysis buffer and beads extracted important community members from biofilms with minimal plant DNA co-extraction (23). The lysate was incubated for 60 min at 65°C, placed in a NucleoSpin Filter tube, and centrifuged for 2 min at 11000x*g* following manufacturer instructions. The filtrate was added to 1.6 ml of Buffer PC and extraction continued following recommended protocol steps. For tropical drought, DNA extractions were performed using the QIAGEN RNeasy Power Water Kit, which extracts total RNA and DNA. This method utilizes the same principle used in temperate grasses in that vortexing whole leaves with beads and lysis buffer increases bacterial DNA extractions while minimizing plant DNA co-extraction. Instead of a membrane filter, a single leaf (∼0.5-2 g) was used for extraction following the manufacturer’s protocol to isolate DNA. All final products were stored at -80°C.

Samples underwent a two-step PCR amplification to attach Illumina adaptor sequences and barcodes as detailed in Supplemental Materials. The first PCR step used chloroplast excluding primers 799F and 1115R targeting the V5-V6 region of the 16S rRNA gene (24). The product from the first PCR was used as template for the second PCR to attach unique Access Array Barcodes (Fluidigm, San Francisco, CA, USA) using previously published methods (42). The amplicons were pooled and sequenced on Illumina MiSeq Platform, with 251 bp paired-end sequencing chemistry. Illumina PhiX was spiked-in (∼ 25%) to account for the low base diversity. Sequencing was performed at the Genomics Resource Laboratory (University of Massachusetts-Amherst).

### Sequence Analysis

Using the QIIME2 (43) pipeline, paired-end reads were demultiplexed, merged, trimmed to 315 bp, and binned using DADA2 (43) inferring amplicon sequence variants (ASVs). Taxonomic identities were assigned using the naïve Bayes sklearn classifier trained with the 799F/1115R region of the Greengenes 13_8 database.

In temperate grass species, 2,142,917 high quality paired-end reads were kept while 560,063 (20.7%) reads failed quality control and were removed. After filtering out chloroplast and mitochondria sequences, 1,853,078 reads were left (4,340-75,714 reads per sample) (Supplemental Table 2). Seven samples were removed from analysis due to inadequate coverage, leaving 36,277 ASVs representing 39 different phyla.

In tropical grass species, 3,604,936 high quality paired-end reads were kept while 1,555,207 (30.1%) reads failed quality control and were removed. After filtering out chloroplast and mitochondria sequences, 3,522,914 reads remained (10,478-116,876 reads per sample) (Supplemental Table 3). Two samples were removed from analysis due to inadequate coverage, resulting in 6,400 ASVs representing 28 different phyla. All tropical and temperate samples were rarefied to 4000 reads which sufficiently captured the diversity in both tropical and temperate systems.

### Statistical Analyses

Because of the differences in experimental procedures, all analyses were performed separately on samples from tropical and temperate grasses, and models were never made combining the data. Alpha diversity was calculated using the Shannon Diversity Index, Chao1, Pielou’s Evenness, and ASV Richness. Differences in each alpha diversity metric as a result of species, treatment, and time were calculated using Kruskal-Wallis pairwise comparisons in Qiime2. We additionally assessed changes in Shannon Diversity Index as a result of drought and host species at the end of the experimental period using generalized linear models (GLMs) in R (44). To then understand these changes over time, we used generalized linear mixed models (GLMMs) with plant ID included as a random effect to account for the repeated measures over the course of the experiment (lme4 package (45)). For both GLMs and GLMMs a gamma distribution with a log link was used because alpha diversity is a continuous variable bounded at zero. Correlation of variables was tested using the Car package (46) and the best model was selected using AICcmodavg (47). Differences in alpha diversity metrics Chao1, Pielou’s Evenness, and ASV Richness as a result of species, treatment, and time were calculated using Kruskal-Wallis pairwise comparisons in Qiime2. Alpha diversity was assessed separately for the temperate and tropical grasses.

Beta diversity was assessed using weighted UniFrac and Bray-Curtis distances as implemented in Qiime2. Permutational multivariate analysis of variance (PERMANOVA) using the adonis function in the vegan package (48) was used to determine the influence of host species, watering condition, and time on phyllosphere community structure. Post-hoc analysis for pairwise multilevel comparisons was performed using pairwise_Adonis (49). Separate analyses were run for tropical and temperate grass hosts. Results were visualized by creating nonmetric multidimensional scaling (NMDS) plots using vegan and ggplot2 (50). Plant health traits were correlated to community structure and visually overlaid in the NMDS plots using envifit function in the vegan package. Differences in beta diversity were further assessed by identifying bacterial indicator genera for each treatment. Indicators were determined individually for the tropical and temperate grasses using the indicspecies package (51) and visualized by creating ternary plots using the ggtern package (52).

### 16S rRNA Gene Copy Normalization

Because there is a large range of 16S rRNA gene copy numbers between microbial species, we additionally normalized our data based on 16S rRNA gene copy number to determine if this influenced our results (53). 16S rRNA gene copy number was normalized using the q2-gcn-norm plugin in QIIME2. After normalization samples were rarefied to 2000 reads to account for the changes in read number, allowing incorporation of maximum samples and sufficient depth to capture community diversity. All alpha and beta diversity analyses were repeated using gene copy number normalized community data.

### Data Availability

The 16S rRNA gene sequences were deposited in the NCBI Sequence Read Archive (SRA) under BioProject ID PRJNA727243 for tropical grasses and PRJNA727101 for temperate hosts.

Reviewer links:

https://dataview.ncbi.nlm.nih.gov/object/PRJNA727243?reviewer=fi8rccbf5rfo30sdkrcsgtcrhg

https://dataview.ncbi.nlm.nih.gov/object/PRJNA727101?reviewer=va5qcocro5c62sjnl1vea48up4

## RESULTS

### Alpha and Beta Diversity

To characterize the response of grassland phyllospheres to drought we grew two species of temperate grass (Tall Fescue and Orchardgrass) and three species of tropical grass (*Brachiaria brizantha*, *Brachiaria decumbens*, and Brachiaria hybrid) under three different watering conditions: well-watered control, mild drought, and severe drought. Because there is a large range of 16S rRNA gene copy number between microbial species, all analyses were conducted using both gene copy number normalized and non-normalized data. No major differences were found between the two methods, so all data reported directly in the paper are from non-normalized data, but corresponding analyses can be found in the supplemental information (Supplemental Table S4, Supplemental Fig. S3-4).

#### Temperate Grass

Phyllosphere communities on selected temperate grass species were altered as a result of severe drought but the degree to which they changed was dependent on host species. Assessment of alpha diversity using Shannon Diversity Index, Chao 1, Pielou’s Evenness, and ASV richness revealed similar trends regardless of the alpha diversity metric used (Supplemental Figure 5).

Because Shannon Diversity Index incorporates both richness and evenness, we decided to use it to model how alpha diversity was impacted over time by drought severity and host species. We fit 20 different GLMMs, which included possible combinations of host species, leaf relative water content, sampling day, and watering treatment as predictor variables and with host ID as a random effect. The best model, selected using the Akaike information criterion (54), was an interactive GLMM comparing host species and leaf relative water content indicating that alpha diversity was impacted by host species identity and decreasing relative water content, but did not change over time (Supplemental Table S5). To further understand the differences, we built a GLM to look at the combined impact of drought and host species on the last sampling day in order to understand the impact of drought when the drought effect was strongest. This model revealed a significant decrease in alpha diversity as a result of severe drought (p =0.01) but not from mild drought (p=0.6) and that alpha diversity was more impacted by severe drought on Tall Fescue than on Orchardgrass (Fig. 1A).

**Figure 1.**
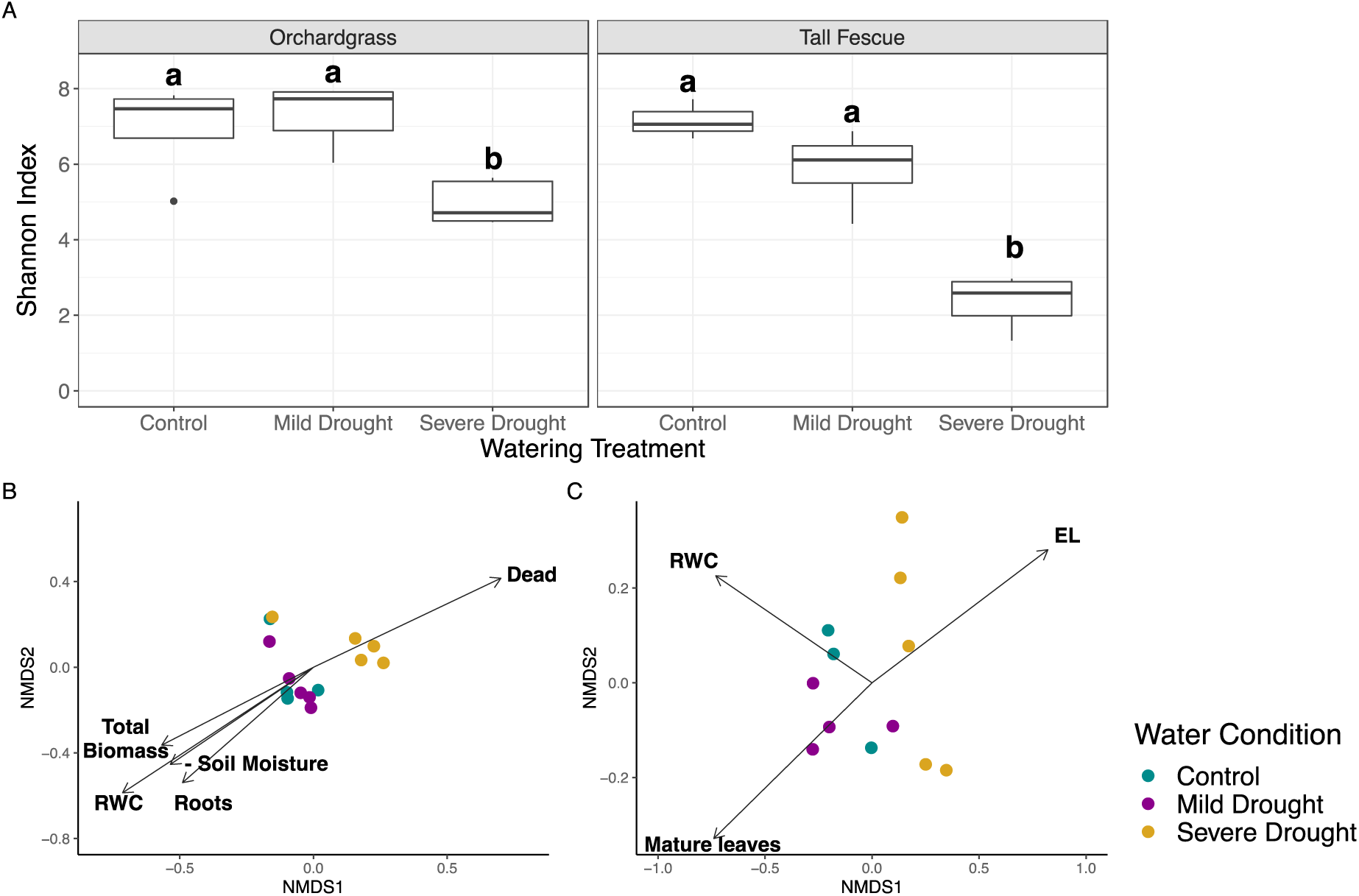
(A) Phyllosphere community diversity (alpha-diversity) was significantly impacted by severe drought and host species on the temperate grass hosts. The diversity measure is based on the Shannon Diversity Index accounting for both abundance and evenness of the species present. Significant differences between treatments within a host species are indicated by different letters above each boxplot. Changes in bacterial community structure correlate with changes in plant traits on (B) Orchardgrass and (C) Tall Fescue. Ordination was calculated using weighted UniFac Distance on samples collected on the last day of drought treatment and visualized by NMDS. Arrows represent significant correlation (P<0.05). Mature leaves, dead leaves, and roots represent the ratio of total biomass. RWC stands for leaf relative water content, EL for electrolyte leakage.

The influence of host species and watering condition on phyllosphere community structure over time was determined using permutational multivariate analysis of variance (PERMANOVA) on weighted UniFrac and Bray-Curtis distances. UniFrac metrics incorporate information on relative relatedness of community members based on phylogenetic distances. We did not observe strong differences between analyses that used weighted UniFrac or Bray-Curtis distances. Results from the PERMANOVA revealed sample day as the strongest driver of community structure and changes in rare taxa (Table 1A). Host species and watering treatment were also significant drivers though each explained a smaller amount of variability (Table 1A). We performed NMDS ordination of weighted UniFrac and Bray-Curtis distances to visualize differences in community structure on the last day of the drought (Supplemental Figure 6). Bacterial community structures were correlated to different measured plant traits for each host species. On Orchardgrass, changes in community structure were correlated with higher relative water content, higher overall biomass, and a higher proportion of root mass in the control and mild drought groups while severe drought had higher proportions of dead leaves (Fig. 1B). On Tall Fescue, community structures of control communities were correlated with higher RWC, mild drought communities with higher proportion of mature leaf mass, and severe drought showed increased electrolyte leakage indicating increased level of stress induced injury to plant tissue (Fig. 1C).

**Table 1.** Compositional dissimilarities between bacterial community structures on the temperate grasses were explained by host species, watering treatment, sampling day, and their interactions using a PERMANOVA on weighted UniFrac and on Bray-Curtis distance measures. Because of the significant three-way interactions, independent PERMANOVAs were conducted for each watering treatment and sampling day to better understand the impact of host species, drought treatment, and time on community structure.

| Temperate Host Species |  |  |  |  |  |  |
| --- | --- | --- | --- | --- | --- | --- |
| Potential Drivers | Weighted UniFrac |  |  | Bray-Curtis |  |  |
|  | F | R<sup>2</sup> | P value | F | R<sup>2</sup> | P value |
| Species | 2.512 | 0.025 | 0.003** | 1.825 | 0.020 | 0.008** |
| **Day** | **10.260** | **0.154** | **0.001\*\*\*** | **4.629** | **0.114** | **0.001\*\*\*** |
| Treatment | 2.268 | 0.022 | 0.012* | 1.910 | 0.031 | 0.001*** |
| Species:Day | 0.719 | 0.021 | 0.890 | 1.048 | 0.026 | 0.353 |
| Species:Treatment | 1.111 | 0.022 | 0.299 | 0.996 | 0.016 | 0.469 |
| Day:Treatment | 1.195 | 0.035 | 0.172 | 1.459 | 0.073 | 0.001*** |
| Species:Day:Treatment | 1.371 | 0.082 | 0.036* | 1.285 | 0.063 | 0.012* |

| Host Species | Sampling Day | Weighted UniFrac |  |  | Bray-Curtis |  |  |
| --- | --- | --- | --- | --- | --- | --- | --- |
|  |  | F value | R<sup>2</sup> | P-adj | F value | R<sup>2</sup> | P-adj |
| Orchardgrass | 1 | 0.540 | 0.107 | 0.941 | 0.952 | 0.175 | 0.008** |
|  | **14** | **2.036** | **0.270** | **0.047\*** | **1.369** | **0.199** | **0.001\*\*\*** |
|  | 23 | 2.548 | 0.316 | 0.029* | 1.455 | 0.209 | 0.001** |
|  | 36 | 2.688 | 0.328 | 0.014* | 2.030 | 0.270 | 0.353 |
| Tall Fescue | 1 | 0.943 | 0.158 | 0.528 | 1.037 | 0.172 | 0.469 |
|  | 14 | 1.097 | 0.166 | 0.331 | 1.175 | 0.176 | 0.001** |
|  | 23 | 1.282 | 0.221 | 0.232 | 1.090 | 0.195 | 0.012* |
|  | **36** | **4.273** | **0.487** | **0.003\*** | **2.735** | **0.378** | **0.008\*\*** |

To determine when in the drought period bacterial communities from each host species were significantly affected, we conducted PERMANOVA analyses on each host species to understand how microbial communities were impacted by drought on each sampling day. By looking at each individual host species, we can better understand how susceptible to drought each community is based on its host species. Changes in community structure were first observed on Orchardgrass on the second day of sampling, day 14, but were not seen until the last day of sampling, day 36, on Tall Fescue (Table 1B) (Supplemental Figures 7-8).

#### Tropical Grass

Phyllosphere communities from all tropical grasses were altered as a result of severe drought but no statistical differences were detected between host species. To determine which variable affects phyllosphere communities we conducted the same modelling process used on the temperate grass hosts. Each of the alpha diversity metrics resulted in similar trends, so Shannon Diversity index was chosen for the modelling process (Supplemental Figure 8). The best model was a GLMM comparing a single categorical fixed effect, watering treatment, with host ID as a random effect (Supplemental Table S6). This model indicates that alpha diversity significantly decreased as a result of severe drought (p=0.002), but that mild drought and time had no significant impact. To further understand the effect of drought, we built a GLM to look at the impact of drought and host species on the last sampling day. This model revealed a significant decrease in alpha diversity as a result of severe drought compared to the control treatment (p<0.001), but showed no significant difference between host species (Fig. 2A).

**Figure 2.**
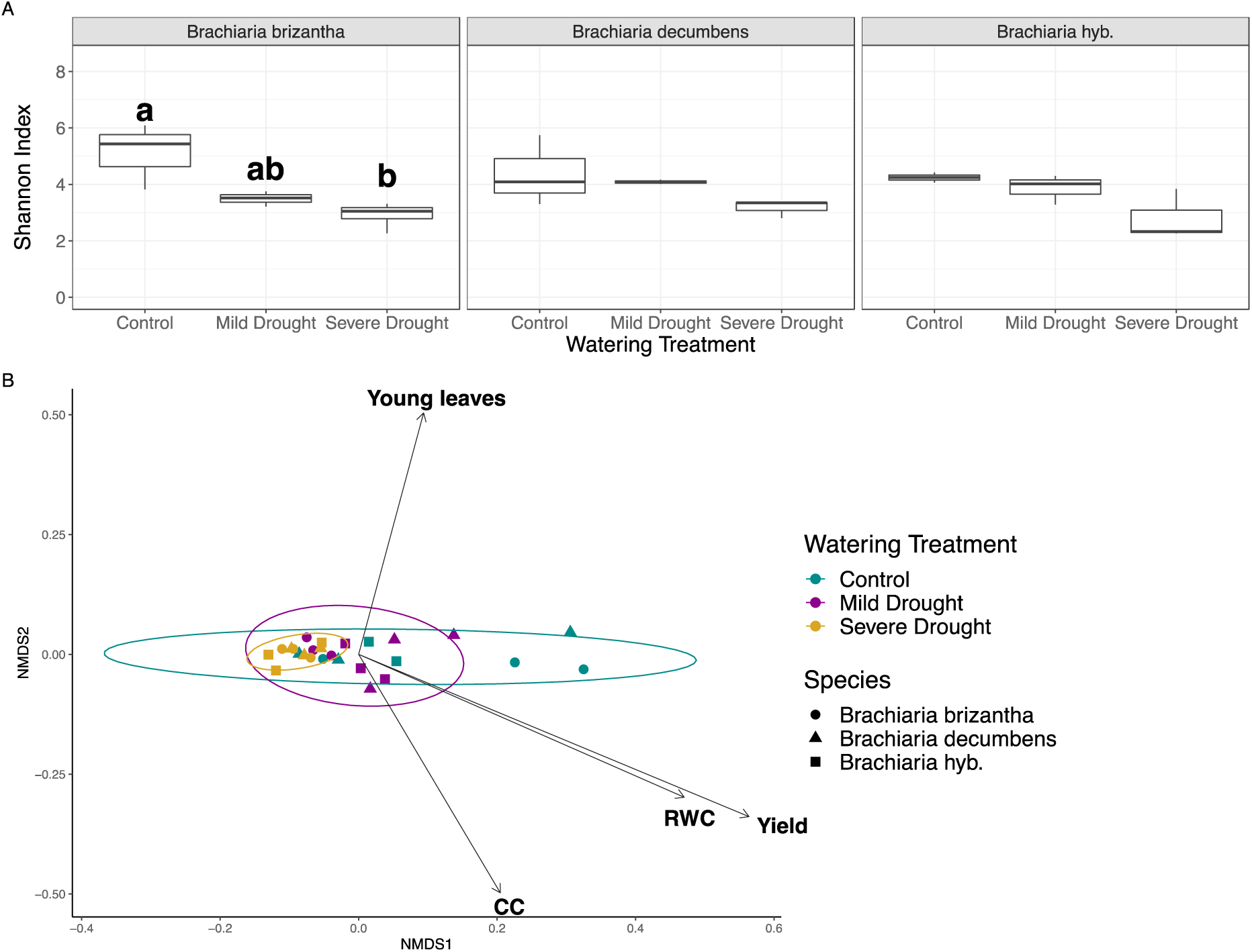
(A) Phyllosphere community diversity (alpha-diversity) was significantly impacted by severe drought, but showed no difference between host species on the tropical grass hosts. The diversity measure is based on the Shannon Diversity Index accounting for both abundance and evenness of the species present. Significant differences between treatments within a host species are indicated by different letters above each boxplot. (B) Bacterial communities from control and mild drought treatments correlated with plant traits on tropical grass species. The NMDS ordination was calculated using weighted UniFrac Distance of community samples collected on the last day of drought treatment. Arrows represent significant correlation (P<0.05). RWC stands for leaf relative water content, CC for chlorophyll content measured by chlorophyll extraction, and yield for photochemical efficiency.

The influence that host species and watering condition had over time on beta diversity was determined using PERMANOVA. Results from the PERMANOVA using weighted UniFrac and Bray-Curtis distances revealed watering treatment as the only significant predictor of community structure (Table 2). Post hoc analysis on weighted UniFrac anlayses revealed, only severe drought was significantly different from the control (R^2^=0.297, p=0.022) and from the mild drought communities (R^2^=0.351, p=0.001), but that control and mild drought treatments were not different from each other (R^2^=0.094, p=0.196). Performing an NMDS ordination of the weighted UniFrac and Bray-Curtis distances allowed us to visualize differences in community structure on the last day of the drought. Because no significant difference between host species communities were observed, NMDS was plotted together. When community structure was correlated to measured plant traits, communities under control and mild drought conditions were associated with higher chlorophyll content and photochemical efficiency (yield), higher relative water content, and a higher proportion of young leaves (Fig. 2B) (Supplemental Figure S10).

**Table 2.** Drought treatment was the strongest driver of community structure in tropical grass species. Compositional dissimilarities between bacterial community structures on the tropical grasses were explained by host species, watering treatment, sampling day, and their interactions using a PERMANOVA on weighted UniFrac and on Bray-Curtis distance measures.

| Tropical Host Species |  |  |  |  |  |  |
| --- | --- | --- | --- | --- | --- | --- |
| Potential Drivers | Weighted UniFrac |  |  | Bray-Curtis |  |  |
|  | F | R<sup>2</sup> | P value | F | R<sup>2</sup> | P value |
| Species | 0.558 | 0.013 | 0.775 | 1.342 | 0.034 | 0.180 |
| Day | 1.887 | 0.046 | 0.087 | 1.454 | 0.037 | 0.131 |
| Treatment | **4.198** | **0.103** | **0.003\*\*** | **1.956** | **0.049** | **0.029\*** |
| Species:Day | 0.740 | 0.036 | 0.675 | 0.583 | 0.030 | 0.952 |
| Species:Treatment | 0.856 | 0.042 | 0.548 | 1.191 | 0.061 | 0.252 |
| Day:Treatment | 1.215 | 0.059 | 0.268 | 1.435 | 0.073 | 0.091 |
| Species:Day:Treatment | 0.858 | 0.084 | 0.637 | 0.779 | 0.079 | 0.860 |

### Taxonomic Analysis

To gain a deeper understanding of community changes seen between watering conditions at the end of the drought, we compared changes in taxonomy to observed changes in community structure. Using NMDS ordination of Bray-Curtis and weighted UniFrac distances (Supplemental Fig. S11) and taxa barplots comparing relative abundance (ANOVA with Tukey’s test post hoc analysis), *Gammaproteobacteria* significantly increased under severe drought in all six host species (Fig. 3) (Supplemental Table 7). Next, we compared changes in dominant genera between treatments. In temperate grasses, a significant decrease was observed in *Bacillus* and *Pseudomonas* between control and severe drought conditions (Fig. 4A). The abundancies of *Acinetobacter, Bacillus, Sphingomonas,* and *Staphylococcus* all showed an increasing trend in mild drought and a significant decrease in severe drought communities. *Buchnera* significantly increased and four genera (*Cornyebacterium, Delftia, Erwinia, Streptococcus*) remained unchanged between watering conditions. In the phyllosphere of tropical grasses, six taxonomic groups showed no difference, while *Methylobacterium, Sphingomonas,* and *Hymenobacter* significantly decreased under mild and severe drought, and *Deinococcus* under severe drought (Fig. 4B). In order to understand how changes in relative abundance related to bacterial load on plant leaves, cell counts per leaf area were conducted. In temperate grasses, a decreasing but not significant trend was observed under severe drought conditions (Fig. 5A). For tropical grasses, all three species showed a significant decrease in cell counts on hosts under severe drought compared to both control and mild drought conditions (Fig. 5B).

**Figure 3.**
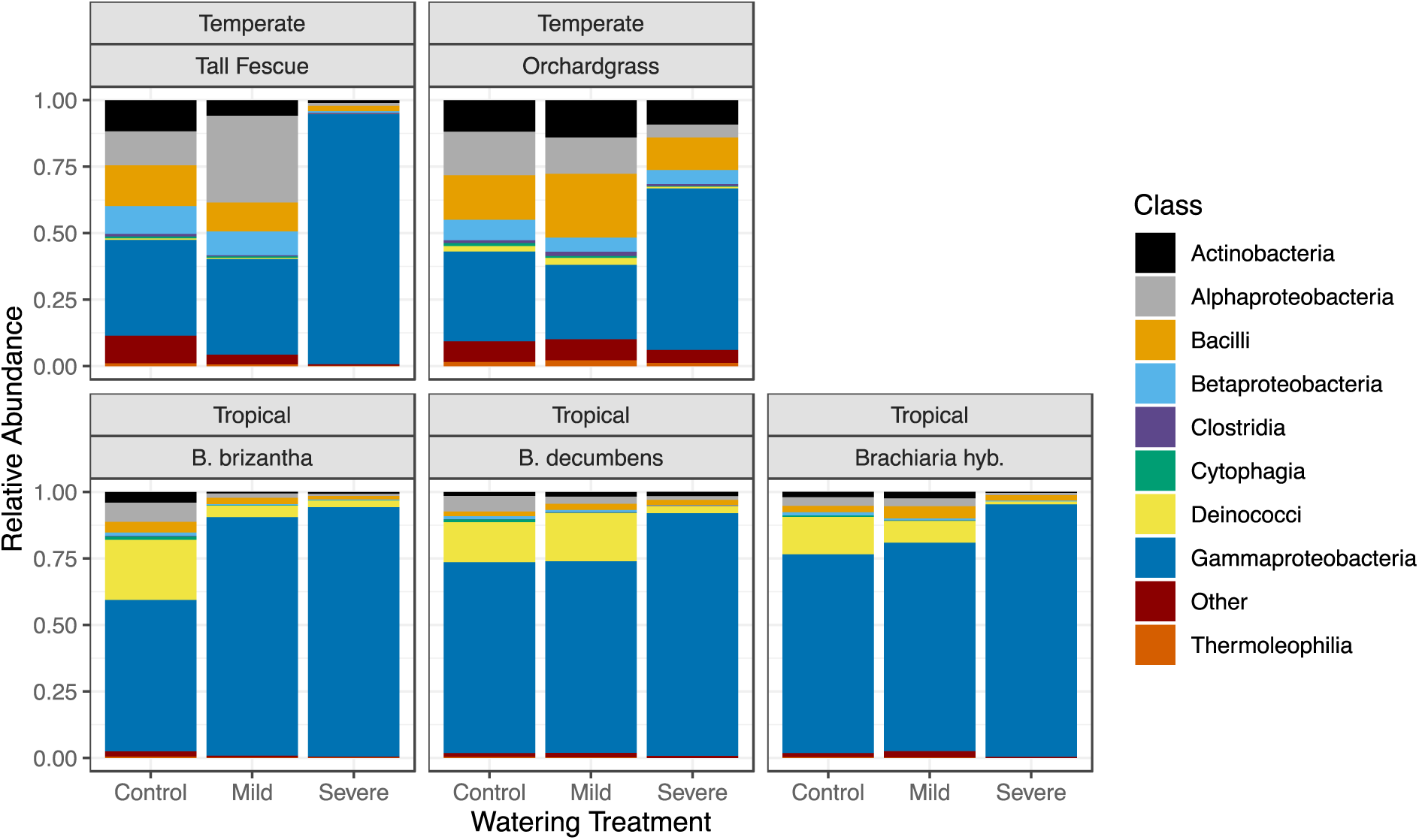
Average relative abundance of the dominant class of phyllosphere bacteria collected at the end of the experimental period. Each treatment was calculated based on 5 biological replicates in the temperate grass species and 3 biological replicates in the tropical grass species. Changes in bacterial classes by watering treatment were tested using an ANOVA followed by a *post hoc* Tukey multiple comparison test (Supplemental Table S7). Gammaproteobacteria significantly increased as a result of severe drought on Tall Fescue (p.adj=0.0002), Orchardgrass (p.adj=0.02), and on the tropical grass species (p.adj=0.002).

**Figure 4.**
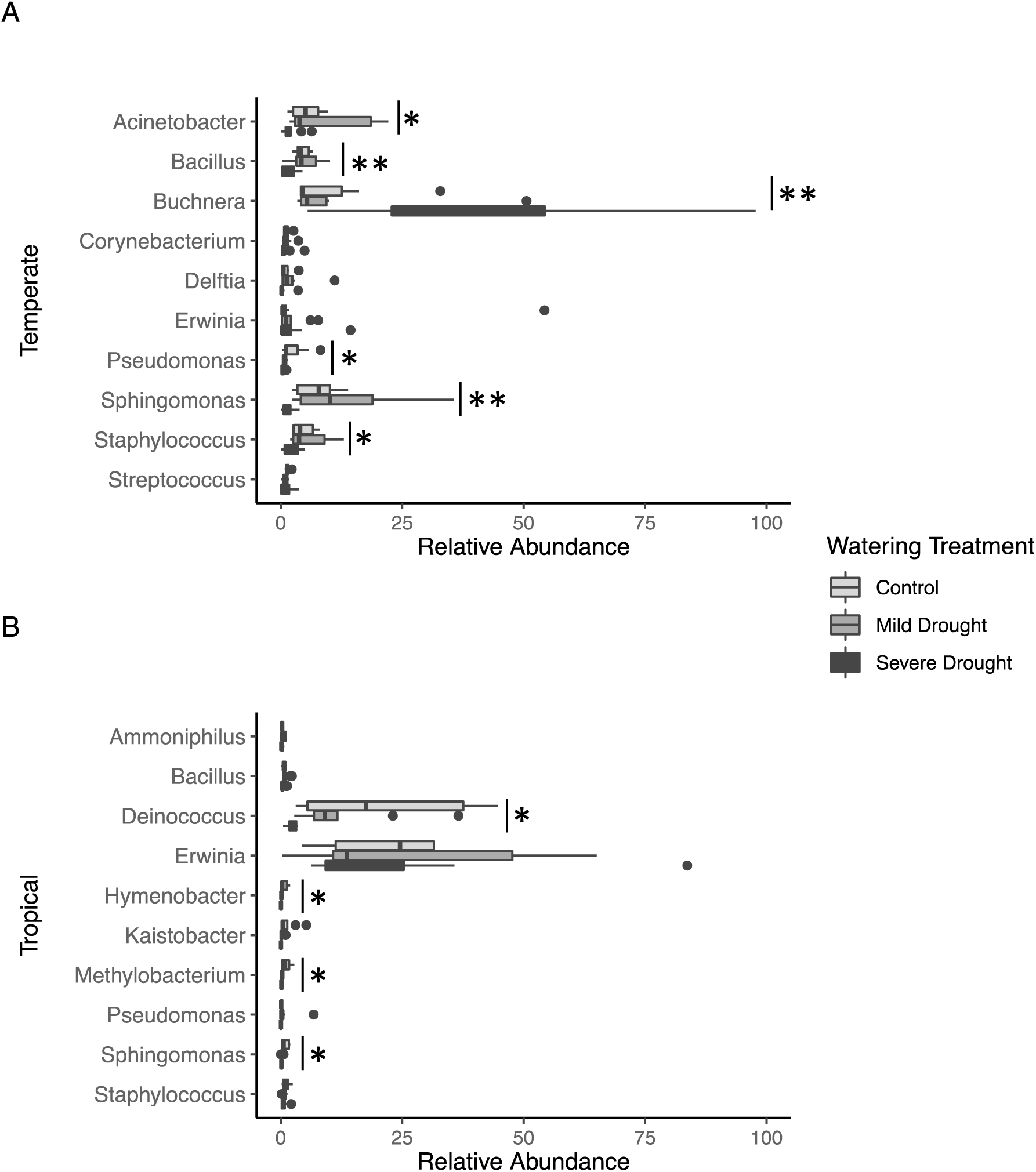
Average relative abundance of the top 10 most dominant genera in bacterial communities from (A) Temperate grass species and (B) Tropical grass species. In temperate grass species each treatment is represented by 10 biological replicates, and in tropical species by 9 biological replicates. Differences were calculated using analysis of variance (ANOVA) followed by a *post hoc* Tukey multiple comparison test. Data are displayed using boxplots, which visualizes the distribution of the data. The box represents the interquartile range with the line in the middle representing the mean value of the data. The whiskers coming from the sides of the boxes represent the minimum and maximum quartiles, and the circles represent outliers. Additionally, significance levels are assigned as P>0.05 (not significant); * P ≤ 0.05; ** P ≤ 0.01; *** P ≤ 0.001.

**Figure 5.**
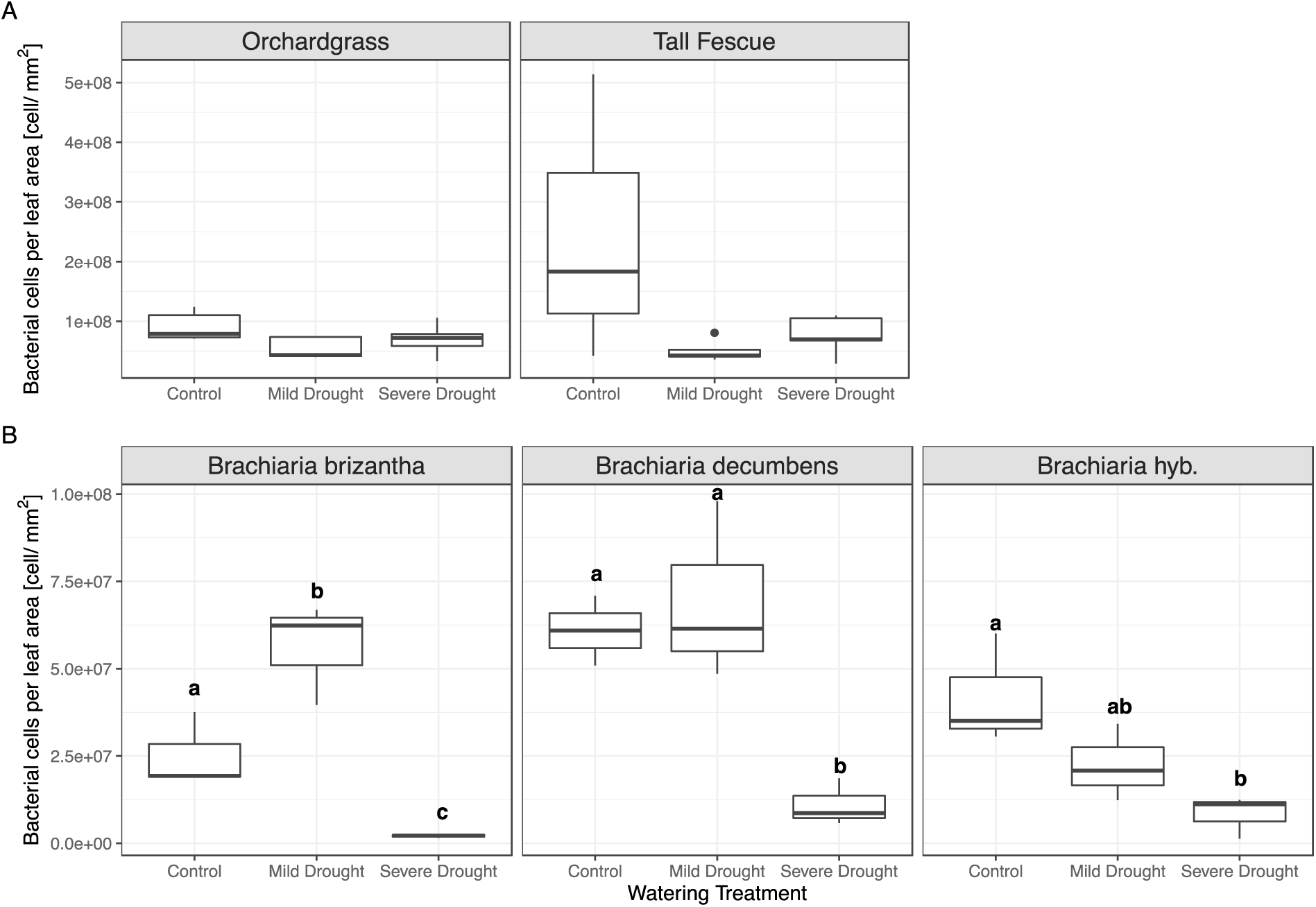
The total number of bacterial cells per leaf area in (A) temperate and (B) tropical grass species. Each watering treatment is represented by 5 biological replicates from the temperate grass hosts and 3 biological replicates for the tropical grass hosts. Number of bacterial cells on tropical grasses decreased significantly as a result of severe drought for each individual species. Significant differences between treatments within a host species are indicated by different letters above each boxplot. Bacterial cells per leaf area (mm^2^) were calculated by washing bacterial cells off the surface of leaves, counted using epifluorescence microscopy, and compared to leaf areas calculated using ImageJ.

### Indicator Analysis

To determine if any phyllosphere bacteria are indicators of watering condition, we performed an indicator analysis (detailed in Supplemental Table S8) (26, 55). In the temperate grass species, 57 genera were indicators for control treatments, 9 were indicators in mild drought, 23 were indicators shared between control and mild drought, and 2 were indicators of severe drought. The relative abundance distribution of genera between treatments was visualized in ternary plots for the 9 dominant phyla (Fig. 6A). Several of the most dominant genera were indicator taxa. *Pseudomonas* was an indicator of the control community; *Sphingomonas* was an indicator of the mild drought community; *Bacillus*, *Hymenobacter*, and *Methylobacterium* were indicators of both control and mild drought communities; and *Buchnera* was an indicator only of severe drought (Table 3).

**Figure 6.**
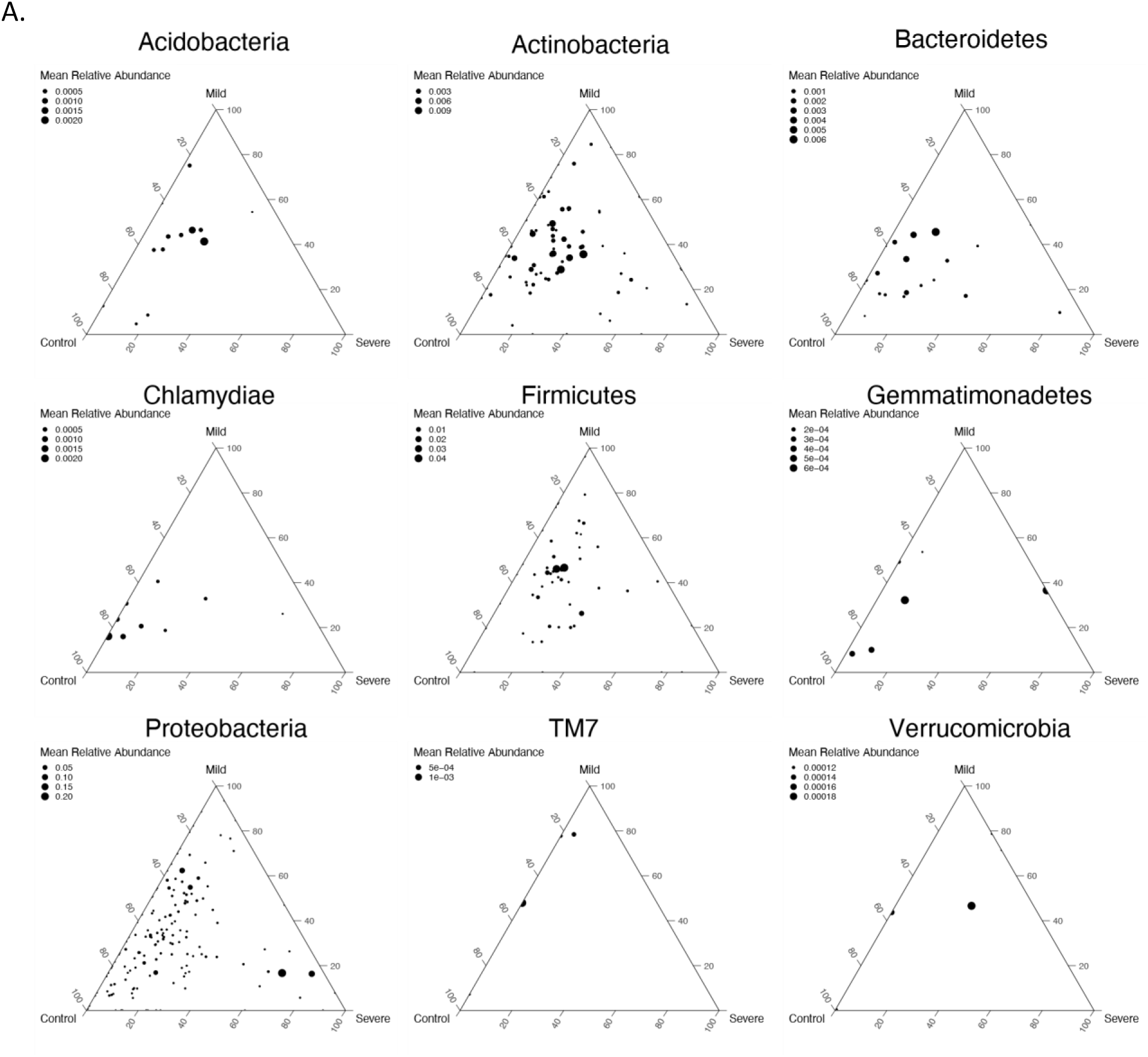
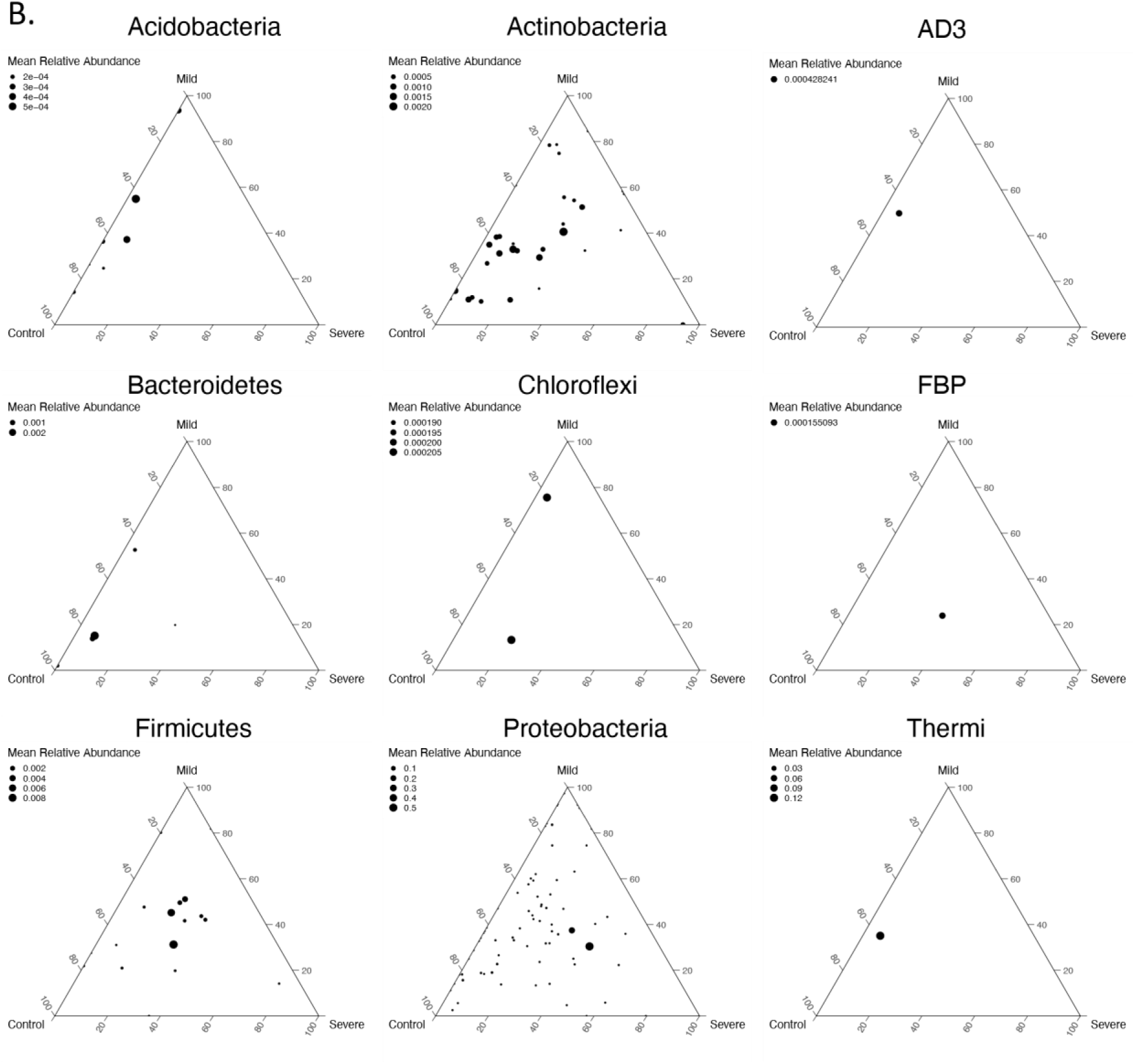
Ternary plots of the relative abundance of genera for the 9 most abundant phyla in each watering condition for bacterial communities from (A) Temperate and B) Tropical grass hosts. Each small circle represents a genus fitted onto an x-y-z coordinate system, where each corner represents one of the three watering conditions (Control, Mild Drought, Severe Drought). Therefore, points closer to the corners have a greater proportion under that watering condition compared to points in the middle which have more equal proportions for each watering condition.

**Table 3.** Functions that have been observed in the dominant genera found on tropical and temperate grass species. These genera were also identified as indicator genera in this study.

| Genus | Indicator | Observed Functions | Source |
| --- | --- | --- | --- |
| *Bacillus* | Temperate<br>control-<br>mild | Biological control agent; in rhizosphere increase biomass and growth, phytohormone production (IAA, GAs, cytokinins); under drought conditions promote plant growth through increased availability of water, nutrients and hormones and regulate drought response genes. | (79–81) |
| *Deinococcus* | Tropical<br>control-<br>mild | Resistant to UV damage and radiation; degradation of aromatic compounds | (74, 83, 84) |
| *Hymenobacter* | Temperate<br>control-<br>mild<br>Tropical<br>control | Potential phyllosphere “hub” taxa; can produce carotenoids which can be involved in photosynthesis, stabilize cellular membranes, and act as antioxidants; and have genes involved in UV-damage DNA repair | (23, 85, 86) |
| *Kaistobacter* | Tropical<br>control | Potential disease suppression; associated with healthy plants in rhizosphere and phyllosphere studies but function unknown | (87–89) |
| *Methylobacterium* | Temperate<br>control-<br>mild<br>Tropical<br>control | Promote plant growth; nitrogen fixation;<br>phytohormone production; protect against<br>desiccation by producing osmo-protectants | (72, 90–<br>92) |
| *Pseudomonas* | Temperate<br>control | Produce phytohormones (IAA, cytokinins, and<br>giberellins), protect against diseases (through<br>antibiosis, niche exclusion, quorum sensing,<br>and induction of systemic resistance)<br>Some pseudomonads are also plant pathogens | (93–95) |
| *Sphingomonas* | Temperate<br>mild<br>Tropical<br>control | Increase water uptake under drought through<br>enhanced root growth, pathogen protection | (15, 23,<br>96) |

In the tropical grasses, 25 indicators were associated with control treatments, 3 with mild drought, 3 for control and mild drought combined, and none for severe drought. The distribution of genera between treatments was visualized in ternary plots for the 9 most dominant phyla (Fig. 6B). Indicator taxa that were also dominant phyllosphere genera were *Hymenobacter*, *Kaistobacter*, *Methylobacterium*, and *Sphingomonas* which were indicators of the control condition and *Deinococcus* which were an indicator of control and mild drought conditions.

## DISCUSSION

This study investigated how phyllosphere microbial communities from five different forage grass species responded to drought. Three species of C_4_ tropical grasses and two species of C_3_ temperate grasses were grown as well-watered controls and under different drought conditions in a greenhouse to investigate how plant taxonomy and traits influence bacterial community assembly and response to stress. Few studies have compared phyllosphere studies in the greenhouse to studies in the field, but they consistently show that bacterial communities in the greenhouse had lower numbers of bacteria, reduced bacterial diversity, and a shift in dominant bacterial class (56, 57). Despite these differences, these studies concluded that greenhouse studies provide an important controlled framework to study phyllosphere community assembly and succession.

Common characteristics in the microbial communities across each grass species were observed, such as high abundance of *Gammaproteobacteria* and *Alphaproteobacteria* consistent with previous grassland phyllosphere studies (18, 23). Other groups, including *Actinobacteria*, *Deinococci, Betaproteobacteria,* and *Bacilli,* were among the most dominant classes of phyllosphere bacteria on every grass species, but occurred at different abundances between host species. Previous work comparing phyllosphere communities from temperate and tropical tree species also found that some phyla were consistently present on all tree species and are likely common phyllosphere residents, indicative of potentially beneficial effects on plant hosts (58). Phyllosphere microbes directly promote plant growth and stress tolerance through phytohormone and nutrient production and protection from UV damage (14, 59, 60). Identifying microbes, both taxonomically and functionally, that can continue to survive and promote plant health under changing climatic conditions will be important for creating resilient agricultural systems.

### Host Species Impact on Community Structure

Consistent with previous studies, we found that plant species affected microbial community structure, but aspects of community structure varied by host species. The microbial communities from the temperate grass species were distinguishable from each other under severe drought conditions while the three tropical grasses were not distinguishable when using weighted UniFrac and Bray-Curtis distances. The tropical grass species are more closely related to each other than the temperate grass species (Supplemental Fig. S12). Phylogenetic relationships of the host could therefore account for the varying effects each plant species has on microbial community structure. Kembel et al. (25) found that different bacterial community structures could be explained by different host attributes including resource uptake strategies, leaf morphology, and physiology. This confirmed our first hypothesis that host identity influences microbial community assembly and that communities coming from closely related plants are more similar to each other.

### Bacterial Stress Response Correlates to Plant Stress Response

Different grass species employed dissimilar strategies for dealing with drought stress. While some universal trends were observed in phyllosphere response to drought, the differences seen in resource uptake strategies between host species under control conditions were exacerbated under drought conditions. Three methods of drought tolerance used by plants are commonly identified: drought escape, drought avoidance, and tolerance of dehydration (61). Drought escape is often seen as dormancy of the plant through dehydration of tissue. Instead of complete dormancy, Orchardgrass and Tall Fescue enter a state of inactivity where no growth occurs and leaves senesce (61). Conversely, tropical grass species remain productive throughout the dry season and previous greenhouse studies that were consistent with ours showed only minor decreases in biomass under drought conditions (62, 63). None of the grasses compared in this project utilize the dormancy strategy, but instead employ some combination of the other two strategies (Supplemental Table S9). Every grass species showed a decrease in phyllosphere alpha-diversity, but the extent varied greatly, with the tropical grass species experiencing a smaller reduction than temperate species. The tropical grasses used in this study each utilize drought avoidance by forming deep-rooting networks to acquire water (Supplemental Table S10). Their roots, reaching two meter depths, have high extraction efficiency allowing continuous water uptake during dry conditions (62, 63). Additionally, because of their continued growth and color retention during low water stress, we propose they exhibit dehydration tolerance. One mechanism the tropical grasses use to tolerate dehydration is stomatal closure at relatively high leaf water potential (62, 64, 65). This is likely the reason that tropical grasses showed a significant decrease in the number of cells present on their leaf while temperate grasses remained unchanged. Since stomata provide an interface for nutrient and water acquisition by the microbes, less plant-provided nutrients and water are available on the leaf surface for the microbes as drought stress increases and plants close their stomata to reduce transpiration (10, 66, 67). Because tropical species have high drought tolerance and employ similar methods to counteract drought stress, we only observed minor changes in their phyllosphere community diversity and structure between treatments.

Conversely, previous work on Orchardgrass and Tall Fescue found the two species have different drought survival strategies which parallels the increased differences observed in their microbial communities (61). Tall Fescue utilized drought avoidance by growing extensive and deep-rooted networks while Orchardgrass relied on water uptake in low soil moisture conditions. Additionally, Orchardgrass shows tolerance of dehydration by protecting the meristem against dehydration and promoting membrane stabilization. In our study, phyllosphere communities found on Orchardgrass showed early changes in community structure as a result of drought, before any plant response was observed. However, by the end of the drought period, we observed a greater decrease in alpha-diversity and community structure was more impacted by severe drought stress even though Tall Fescue plants showed similar signs of stress compared to Orchardgrass (Supplemental Fig. S1). Early response by the Orchardgrass communities could indicate a host-microbe response resulting in increased stress resilience. Overall, changes in diversity as a result of stress were host species dependent, thereby supporting hypothesis 3 and simultaneously raising the question: are phyllosphere communities and the functions they provide a plant trait?

### Drought Stress Changes Phyllosphere Community Diversity and Structure

Under severe drought stress, bacterial communities on leaf surfaces showed some common trends. Confirming hypothesis 2, alpha-diversity decreased, and community structure shifted, resulting in increased relative abundance of *Gammaproteobacteria.* Previous work found that increased phyllosphere bacterial diversity resulted in higher ecosystem productivity, likely relating to the complementarity effect, a situation in which more diverse communities use more available resources as a result of niche partitioning and are therefore more productive (16, 68). Additionally, high diversity frequently results in functional redundancy, which helps promote resiliency and resistance of communities by increasing the likelihood that niches are filled under various environmental conditions (69, 70). Therefore, a loss of bacterial community diversity and change in community structure could indicate decreased plant and bacterial community health.

Potentially beneficial genera found on several grass species were identified as indicators of control or mild drought conditions and were suppressed under severe drought (Table 3). Of particular interest were the indicators of mild drought, either singly or in combination with control, because these are bacteria able to withstand some level of climate stress and could therefore be good biofertilizer targets. The dominant mild drought indicators include *Bacillus, Deinococcus, Hymenobacter, Methylobacterium, Pseudomonas,* and *Sphingomonas* (Table 3). While their interaction with plants in this study is unknown, previous work has shown several different beneficial relationships with plant hosts. Soybean plants inoculated with *Sphingomonas* demonstrated increased growth and drought tolerance (71), while *Methylobacterium* residing in the phyllosphere have been linked to nitrogen-fixation and biomass production (66, 72). Additionally, some *Methylobacterium* and *Deinococcus* are resistant to UV radiation which gives them a selective advantage on the leaf surface and also helps to protect the plant from UV damage (73, 74)

In drought stressed temperate and tropical grasses, the dominant *Gammaproteobacteria* included bacteria from genera known to be potential plant pathogens, such as *Erwinia* (23, 75). These results are consistent with previous research looking at grasslands under climate stress. Aydogan et al. (2018) observed that with the rise in *Gammaproteobacteria,* caused by elevated ambient temperatures in grasslands, the appearance and growing preeminence of potential plant and human pathogens also increased (23).

Temperate grass species also saw changes in the genus *Buchnera*, for which high abundance was an indicator genus of the severe drought treatment (Supplemental Table S5A). *Buchnera,* an aphid endosymbiont (76), has previously been found in grassland phyllospheres under climate stress (23). Throughout our study, aphids were found on the temperate study plants but the higher abundance of *Buchnera* on severe drought plants is consistent with previous studies looking at the effects of drought or temperature stress on plants, which found they are more susceptible to aphid infestation as a result of changes in secondary metabolite production (77, 78). However, we cannot discount the idea that increased pest presence under stress conditions results from changing phyllosphere communities (23). By increasing abundance of potential pathogens and decreasing abundance of commensals and bacterial diversity, drought stress weakened the phyllosphere’s ability to promote plant health and growth.

### Conclusion

Plant productivity is directly influenced by microbes in the phyllosphere, but what factors govern these symbiotic relationships is still unknown. To date, this is the first study to show the significant effects drought has on phyllosphere bacterial communities of grasses. Five commonly used grass species, specifically selected for their variation in native climate zone, taxonomy, and pathways of carbon fixation, were used in order to understand how grassland phyllosphere communities are composed and whether community response to stress is similar. Some similarities in bacterial communities and their response to stress were observed, and the overall trends may be representative for other temperate and tropical grasslands. Bacterial community abundance and diversity decreased while relative abundance of potentially pathogenic bacteria increased, indicating reduced bacterial community health. Additionally, differences in microbial communities in relation to plant traits became more defined under stress conditions. Still, the question remains if bacterial communities can act as a stress response trait for the host plants. Finally, we found that some bacteria indicators of mild drought conditions are potential plant symbionts and are therefore useful targets for biofertilizers designed to promote agricultural systems under climate stress. Future studies should focus on functional interactions between communities and plant host in response to stress conditions in order to understand how bacteria promote plant growth and stress tolerance. By understanding microbe-microbe and plant-microbe interactions, we can better support agricultural practices and discover ways to promote ecosystem productivity through the use of bacteria.

## SUPPLEMENTAL MATERIAL

Supplemental material is available online only.

Supplemental File with methods, figures, and tables, PDF file, 0.7 MB.

## ACKNOWLEDGMENTS

We would like to thank Chris Joyner and the University of Massachusetts College of Natural Sciences Greenhouse Staff for helping to maintain the plants, Michelle DaCosta and Jefferson Lu for guidance and equipment for measuring plant health. We are thankful for financial support from the Lotta M. Crabtree Foundation, and the National Science Foundation – Dimensions of Biodiversity (DEB 1442183).

